# Phenomenon of music-induced opening of the blood-brain barrier in healthy mice

**DOI:** 10.1101/2020.10.03.324699

**Authors:** O. Semyachkina-Glushkovskaya, A. Esmat, D. Bragin, O. Bragina, A. A. Shirokov, N. Navolokin, Y. Yang, A. Abdurashitov, A. Khorovodov, A. Terskov, M. Klimova, A. Mamedova, I. Fedosov, V. Tuchin, J. Kurths

## Abstract

Music plays a more important role in our life than just being an entertainment. It is an even anti-anxiety therapy of human and animals. However, the unsafe listening of loud music triggers hearing loss in millions of young people and professional musicians (rock, jazz, and symphony orchestra) due to exposure to damaging levels of sound using personal audio devices or at noisy entertainment venues including nightclubs, discotheques, bars, and concerts. Therefore, it is important to understand how loud music affects us.

In this pioneering study on healthy mice, we discover that loud rock music below the safety threshold causes opening of the blood-brain barrier (OBBB), which plays an important role in protecting the brain from viruses, bacteria and toxins. We clearly demonstrate that listening loud music during 2 hrs in an intermittent adaptive regime is accompanied by delayed (1h after music exposure) and short-lasting (during 1-4 hrs) OBBB to low and high molecular weight compounds without cochlear and brain impairments. We present the systemic and molecular mechanisms responsible for music-induced OBBB. Finally, a revision of our traditional knowledge about the BBB nature and the novel strategies in optimization of sound-mediated methods for brain drug delivery are discussed.

## Introduction

Sounds, like music and noise, are an integral part of our live, and music is an important aspect of sound. We listen to a variety of music such as classical, popular or rock on audio players, radio and TV, or during concerts. Whatever the type, music is comprised of what are known as notes, which are tones of sounds. Thus, the fundamental aspect of music is based on the concept of sound vibration, which due to deep penetration into the brain and body affect individuals’ mood and emotions positively or negatively at both the behavioral and neuronal level [1,2]. The study of the brain bases for musical listening has advanced greatly in the last 30 years [3]. The evidence from basic and clinical neuroscience suggests that listening to music involves many cognitive components with distinct brain substrates [4]. However, little is known how music affects us and what are mechanisms underlying this phenomenon. The World Health Organization estimated that 1.1 billion teenagers and young adults are at risk of developing noise induced hearing loss (NIHL) due to the unsafe listening of loud music using personal audio devices such as smartphones and MP3 players and exposure to damaging levels of sound at noisy entertainment venues including nightclubs, discotheques, bars, pubs and sporting events [5]. For example, sounds at rock concerts routinely reach levels above 100 dB [6–8], which are considered unsafe for any unprotected exposures exceeding 15 min [9,10]. Rock, jazz and symphony orchestra musicians have been found to be at a significant risk of music-based NIHL [11–14].

The blood-labyrinth barrier (BLB) leakage is an important mechanism of NIHL [15]. Anatomically and functionally, BLB is similar to the blood-brain barrier (BBB) and is comprised of the endothelial cells in the strial microvasculature, elaborated the tight and adherens junctions, the pericytes, and the basement membrane controlling the vascular permeability [15, 16]. Loud sound causes a dramatic change in the strial cochlear-vascular unit [17] and an increase in the BLB permeability leads to a number of NIHL [15, 18].

Since the nature of BLB and BBB is similar and they are two keyplayers controlling the vascular permeability [15, 19], it is expected that loud sound will cause an increase permeability of both BLB and BBB. However, there is no information how loud sound affects the BBB integrity.

In this *in vivo* and *ex vivo* experimental study on the healthy mice, we test our hypothesis that loud music below the safety threshold can cause OBBB. We analyze the systemic and molecular mechanisms underlying this phenomenon and discuss advantages and disadvantages of music-induced OBBB.

## Results

### The window of music-induced OBBB

In the first step, we determined the optimal duration/intensity of music exposure for OBBB as well as the time window of OBBB after listening of loud music (the design of our experiments is presented in Figures 1–3 in SI). Song of the Scorpions “Still loving you” was administered for periods up to 0.25 (continues mode), 1 h and 2 h in an intermittent mode: 60 s – sound and 60 s – pause, at intensities ranging up to 70 dB (moderate), 90 dB (loud) and 100 dB (very loud) [20,21]. Using the *in vivo* real-time fluorescent microscopy [22] in awake behavior mice and optical clearing of skull [23], we determined that a stimulus period of 2 h at 90-100 dB produced a robust increase in the BBB permeability to the Evans Blue Albumin Complex (EBAC, 68.5 kDa) that was observed as bright intensity around the cerebral capillaries (Figure 1b). No changes in the BBB permeability were found in intact mice (no music, Figure 1a) or when the stimulus was 70 dB and shorter times of music exposure (0.25 h and 1 h) was used. This method also revealed the delayed OBBB that was only 1-4 h after music effects but not immediately music-off.

**Fig. 1.**
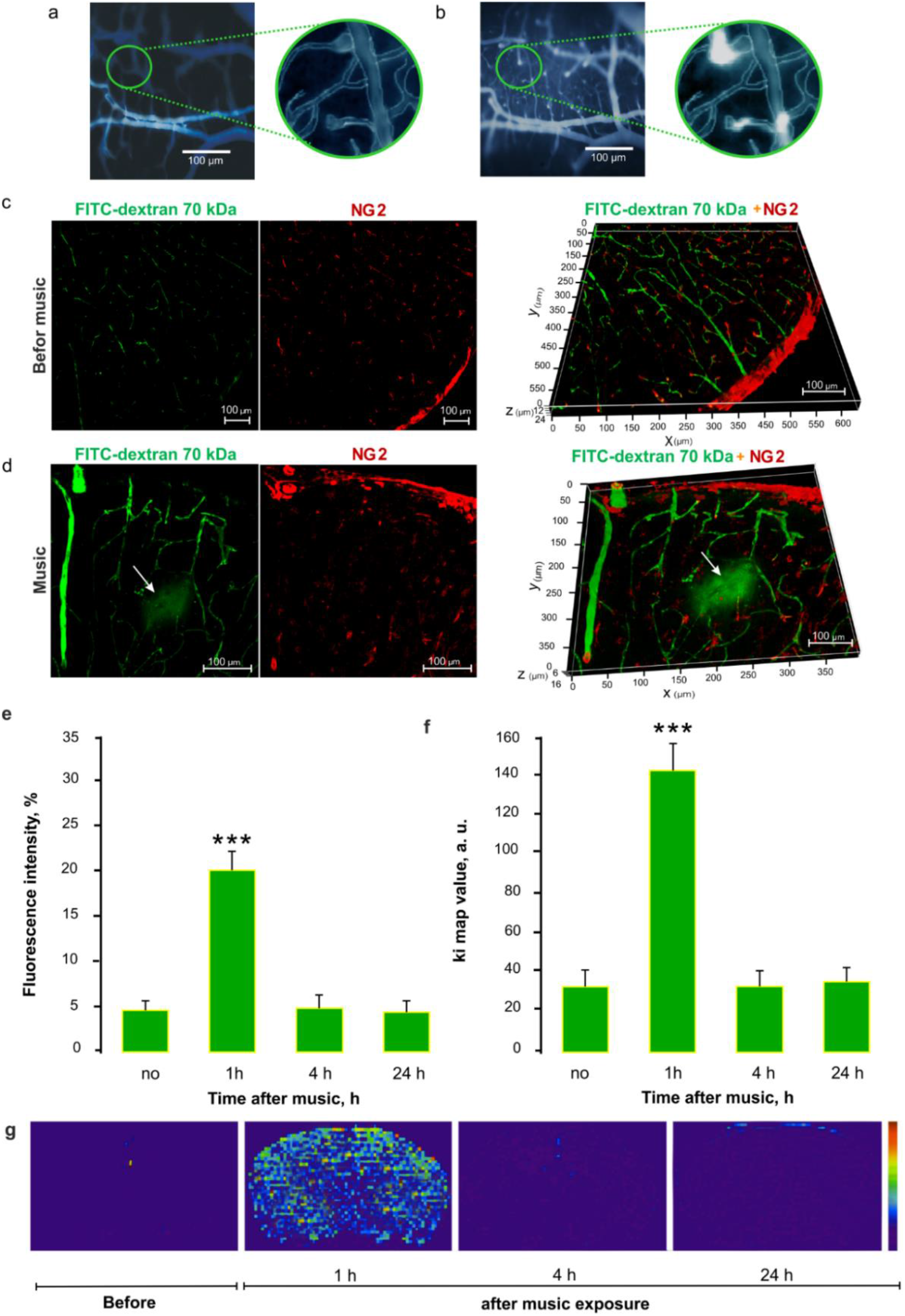
The *ex vivo* and *in vivo* results of music-OBBB. (100 dB, 11-10,000 Hz, 2h intermittent mode: 60 sec – music; 60 sec – pause): a and b - *in vivo* real time fluorescent microscopy of the cerebral microvessels filled ED (no EB leakage) before music exposure (a) and EB extravasation from the cerebral capillaries into the brain tissues 1h after music impact indicating OBBB (b), n=15 in each group; c and d - Confocal imaging of brain slices demonstrating the BBB permeability to FITCD in mice subjected to loud music, where (c) – FITCD intravenous injection but no music exposure (FITCD is constrained to vessels) and d - FITCD injection 1h after music exposure, substantial leakage indicated by diffuse cloud around vessels, n=10 in each group; e - In vivo 2PLSM of the BBB permeability to FITCD in mice subjected to loud music expressed as the percentage fluorescence intensity in perivascular area. Data are presented as mean ± SEM, n = 10, *** - p < 0.01; f and g – The MRI analysis of the BBB permeability to Gd-DTPA in mice subjected to loud music, where f - The Ki values show (arbitrary units) rate of changes in the MRI signal intensity and g - The Ki-maps from the same mice at different time points. Data are presented as mean ± SEM, n = 10, ***p < 0.01.

After the *in vivo* analysis of the BBB permeability to EBAC, all mice were decapitated, their brains were removed and analyzed using spectrofluorometric quantitative assay of EBAC extravasation (Table in SI). These *ex vivo* findings confirmed our *in vivo* data and showed music-induced BBBO in 11 brain regions (Figure 3 in SI). There was no BBBO in the next day after music exposure. Note, that BBBO was observed in all mice (15 of 15) after music exposure (2 hrs) with intensity 100 dB and in 73 % (11 of 15) when music was 90 dB (Table 1 in SI). Since, our interest was to study the effects of very loud music used in nightclubs, discotheques, or rock concerts, where noise levels are recorded to be in excess of 100 dB [14, 24], the next series of experiments were done using the music intensity 100 dB. This choice also was due to the high reproducibility of music (100 dB)-induced OBBB in all mice.

Despite that a quantitative analysis of EBAC is still the most commonly used marker of the BBB integrity, there are some problems arising from its *ex vivo* and *in vivo* application: about 15% of Evans blue dye remains in the endothelium of cerebral vessels, Evans blue dye can be taken by red blood cells and astrocytes [25]. Therefore, for qualitative and quantitative assessment of the BBB integrity we used additional methods for the analysis of the BBB permeability to low and high molecular weight molecules.

In the second step, further optical measurement experiments employed FITC-dextran (70 kDa, hereafter FITCD) injected intravenously 1h after music exposure (the time of OBBB). The BBB disruption was determined both *ex vivo* using confocal microscopy on brain slices and *in vivo* using two-photon laser scanning microscopy (2PLSM) (Fig. 1, c-e). *Ex vivo* confocal microscopy revealed the accumulation of FITCD outside of brain capillaries 1h after music exposure. The leakage of FITCD was visualized by fluorescence outside the vessel walls (Fig. 1d and video 1 in SI). No leakage of FITCD was observed in the control group (no music) (Fig. 1 c and video 2 in SI). The leakage of FITCD in 2PLSM was observed also 1h after music exposure and was quantified by measuring the percentage fluorescence intensity of FITCD in the perivascular area (Fig. 1e). There were no any changes in the BBB permeability to FITCD 4h and 24h after music exposure.

Figures 1 f and g show the post music BBB disruption measured by magnetic resonance imaging (MRI) of gadolinium-diethylene-triamine-pentaacetic acid (Gd-DTPA) leakage. An analysis using rapid T1 weighting for Gd-DTPA is presented in Figure 1g, middle & rightmost images. The rightmost image is significantly brighter uniformly, than the middle one indicating leakage of the tracer molecule. This increase has been quantified by computing the ratio of the 1^st^ and 15^th^ scanned images (Ki map) in Figure 1f, made for times 1, 4, & 24 hrs post music stimulus. The data given in Figures 1f and g indicate a statistically significant (p<0.001) increase in the BBB permeability to Gd-DTPA in all regions of the brain 1h after music exposure. Note, that the Ki map values reached 140±3.7 arbitrary units (p<0.001) only 1h after music exposure, while after 4h and 24h, changes in the Ki values were not observed, compared with the control group (no music).

Altogether, the results of our *in vivo* and *ex vivo* experiments clearly demonstrate that loud music exposure (90-100 dB, 11 – 10.000 Hz, Scorpions “Still loving you”) during 2 hrs in intermittent adaptive mode (1 min – sound; 1 min – pause) is accompanied by delayed (1h after music impact) and short-lasting (during 1-4 hrs) OBBB to low and high molecular weight compounds.

### Mechanisms underlying music-induced OBBB

For both human and mice, music significantly affects emotions and behavior [1,2,26]. There is evidence that mice heard music, which acts as anty-anxiety therapy [26]. However, loud sound triggers stress response in human and animals [5–8,11–18]. Therefore, answered to the question, what is the role of stress in loud music-induced OBBB. The loud sound caused an increase in the plasma level of important stress hormone such as epinephrine up to 3.1-fold (immediately after music) vs. the normal state (9.0±1.5 ng/ml vs. 2.9±0.7 ng/ml, p<0.001, n=10 in each group). One hour after music exposure (the time of OBBB), the level of epinephrine returned almost to the normal value 3.9±1.6 ng/ml (n=10) and was over the control units 4h after music impact (the time of BBB closing) (2.5±0.1 ng/ml, n=10). These data demonstrate that loud music is the stress factor inducing a short increase in the serum epinephrine level. However, the BBB was opened 1h after sound-off, i.e. in the post-stress period when the level of epinephrine was restored.

The rise of epinephrine induced by stress is known to increase in cerebral blood flow (CBF). This can be the initial factor triggering the BBB leakage [27–30]. To test the cerebrovascular changes associated with music-induced OBBB, we measured relative CBF (rCBF) at the venous (the Sagittal sinus) and microcirculatory levels in the same 10 mice before, immediately, 1h, 4h and 24h after sound exposure. Our results demonstrate that immediately after music rCBF was increased in both venous and microcirculatory levels compared with the control group (0.80±0.03 a.u. vs. 0.58±0.01 a.u., p<0.001 for the cerebral microvessels; 1.22±0.01 a.u. vs. 0.83 ±0.02 a.u., p<0.001 for the Sagittal sinus). One hour after sound exposure, when the BBB was opened, rCBF tended to decrease but continued to be higher than the normal level of rCBF (0.77±0.08 a.u. vs. 0.58±0.01 a.u., p<0.05 for the cerebral microvessels; 0.92±0.07 a.u. vs. 0.83±0.02 a.u., p<0.05 for the Sagittal sinus). Four hours after music exposure, when the BBB recovered, rCBF was over the normal units (0.60±0.07 a.u and 0.58±0.01 a.u. for the cerebral microvessels; 0.80±0.06 a.u. and 0.83±0.02 a.u. for the Sagittal sinus, respectively). In the next day, rCBF remained at a normal level (0.56±0.02 a.u and 0.58±0.01 a.u. for the cerebral microvessels; 0.85±0.04 a.u. and 0.83±0.02 a.u. for the Sagittal sinus, respectively).

The tight junction (TJ) proteins such as claudin-5 (CLDN-5), occluding (OCC) and zonula occludens (ZO-1) play a crucial role in regulation of the BBB permeability [19]. Therefore, we studied the integrity of the TJ proteins immediately after music effects, during OBBB (1h after music), and in the time of BBB recovery (4h and 24h after music) compared with the control group (no music). The data of figure 2a show that the signal intensity from CLDN-5 and OCC but not from ZO-1 were significantly decreased in the time of OBBB. There were no changes in the TJ assembly immediately after listening of music as well as in the time of BBB restoration (4h and 24 after music).

**Fig. 2.**
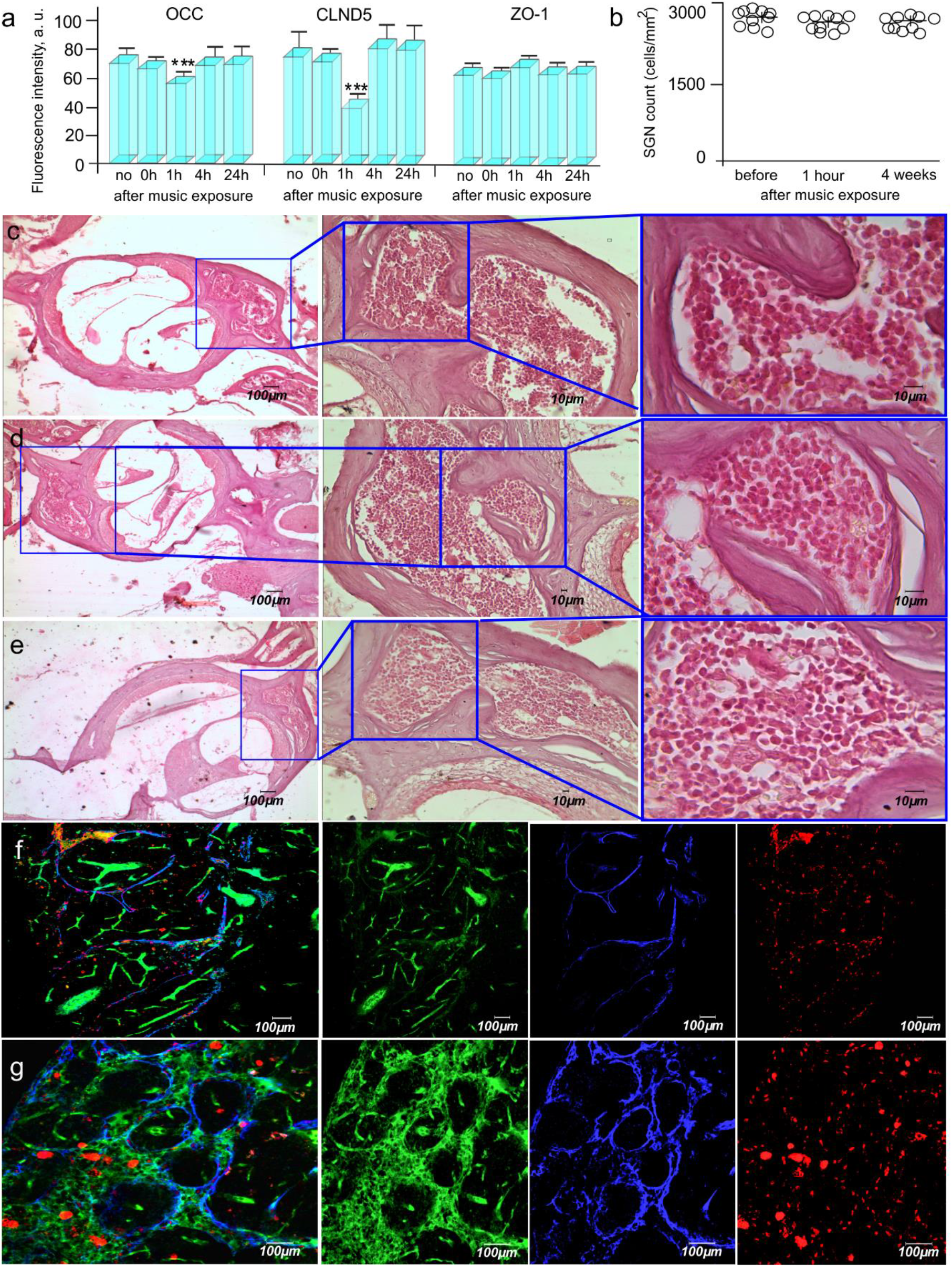
Mechanisms underlying music-induced OBBB: **(a)** The signal intensity from TJ proteins in the control group (before music exposure) and 1-4-24 hrs after music impact (n=10 for each group): *** - p<0.001 vs. the control group; **(b)** quantification of SGN cell density before (no music), 1h and 4 weeks after music exposure; **(c-e)** Cochlear histology: views (64.6x, 246.6x, 774x) of hematoxylin and eosin-stained cochleae of mice before (c), 1h (d) and 4 weeks (e) after music impact; **(f and g)** Confocal imaging of dcLNs in intact mice (f, no music) and in mice with music-OBBB (g, 1h after music exposure): music-OBBB for FITCD was accompanied by lymphatic clearance of FITCD from the brain with its accumulation in the enlarged lymphatic vessels of dcLNs that was not observed in intact mice.

To study the role of the auditory system in OBBB, we studied the effects of loud music on BBB in deafness mice. The results clearly show no sound effects on the BBB permeability to EBAC in mice with hearing loss suggesting the important role of the auditory system in a music-induced OBBB (0.12±0.03 μg/g vs. 0.10±0.08 μg/g, respectively, n=10 in each group)).

To analyze the short-term and delayed effects of loud music on the brain and auditory system injury, whole brain and the cochlear histologic examination was performed using haematoxylin and eosin staining and TUNEL staining for apoptosis 1h (the time of OBBB), 4h (the time of BBB closing), and 4 weeks (delayed effects) after music exposure. The timing of the experiments was dictated by the fact that sound damage may not appear immediately, but after a certain time. For example, peripheral synaptic connections of cochlear neurons are the most vulnerable elements in the cochlea and, in the vast majority of cases, cochlear nerve fibers degenerate only long time after the sound trauma [31]. In animal models exposed to sound stress, hair cell loss can be widespread within hours [31–35], whereas the loss of the spiral ganglion cell (SGCs) is typically not detectable for several weeks to month after sound exposure [31,34,35]. The apoptosis evaluation after ultrasound-OBBB usually performed for several hours to 4-5 weeks after OBBB [36, 37]. Therefore, the time of analysis of SGCs and apoptosis was chosen 1h (the time of OBBB) and 4 weeks (delayed effects) after music exposure.

Figures 2 b-e present cochlear histology and quantification of SGNs density in intact mice (no music), 1h and 4 weeks after music exposure. Our results did not reveal any morphological changes in SGCs after short and delayed effects of loud music on the cochlear system. We found that loud music did not induce apoptosis in the mouse brain tissues (data not presented). A histological analysis of brain tissues and the cerebral vessels also did not show any sound-mediated injuries 1h, 4h and 4 weeks after music exposure (Figure 5 in SI).

In our previous experiments we demonstrated that OBBB by photodynamic [38] or infrasound [39, 40] effects induces the activation of lymphatic clearance of tracers crossed OBBB that is an important mechanism of recovery of the brain [41]. Using confocal imaging of the deep cervical lymph nodes (dcLNs), which are the first anatomical station of the cerebral spinal fluid (CSF) outflow, we clearly uncover that music-induced OBBB for FITCD was accompanied by FITCD lymphatic clearance from the brain with its accumulation in the enlarged lymphatic vessels of dcLNs (22.3±1.5 μm vs 37.3±2.0 μm, p<0.001) that was not observed in intact mice (Figure 2 f and g and video 3 and 4 in SI).

Additionally, to exclude effects of anesthesia (2% isoflurane), which we used in *in vivo* experiments (MRI and 2PLSM), on the BBB permeability, we studied the level of EBAC in the brain tissues in mice 30 min after anesthesia (2% isoflurane). We did not find any difference in the EBAC level in the brain between intact and anesthetized mice (0.17±0.01 μg/g vs. 0.15±0.09 μg/g, respectively). This fact allows us to conclude that short time (30 min) and low dose (2% isoflurane) did not affect the BBB permeability in mice.

## Discussion

Music plays a more important role in our life than just being a source of entertainment. Music is even a powerful therapy that will make calm down both humans and animals, including mice [1,2,26]. However, the unsafe listening of loud music triggers the development of NIHR in millions of people, in particular, in teenagers and professional musicians [5, 8, 11, 13–18]. Therefore, it is important to understand how loud music affects us, especially in nightclubs, discotheques, rock concerts, where the continues sound levels are in excess of 100 dB and are produced for several hours [14, 24].

In this experimental study on healthy mice, for the first time we here demonstrated that the listening of loud rock music (90-100 dB, 11 – 10.000 Hz, Scorpions “Still loving you”) is accompanied by delayed (1h after music exposure) and short-lasting (during 1-4 hrs) OBBB. Music-induced OBBB for high molecular weight molecules such as EBAC 68.5 kDa and FITCD 70 kDa has been clearly shown by *ex vivo* and *in vivo* experiments using conventional fluorescence microscopy and 2PLSM. Moreover, using MRI, have found OBBB for low molecular weight molecules, Gd-DTPA (928 Da). Note, that mice have hearing with a frequency sensitivity 1-90 kHz, i.e. music used in our experiments is audible for them [42].

We have chosen the intermittent music treatment (1 min – sound; 1 min – pause) because the safe listening time for a continues sound of 100 dB is 15 min [24]. However, it has been no effect found on BBB when mice listened continues music of 100 dB during 15 min (Table in SI). Therefore, the longer music exposure during 1h and 2 h in a repetitive mode has been chosen for an adaptation to loud sound. When stimuli are repeated, neural activity is usually reduced due to adaptation [43]. This neural repetition effects have been reported at multiple spatial scales from the level of individual cortical neurons to the level of hemodynamic changes as adaptive mechanisms of the nervous system [44–46]. We have uncovered that a stimulus period of 2 h at 90-100 dB produced significant OBBB. No changes in the BBB permeability have been found when the stimulus was 70 dB or when 1h music exposure was used.

Using a 2h intermittent music exposure, we have found no appearance of apoptotic cells, morphological alterations in the brain tissues, or in SGCs 1h after music impact when the BBB was opened and 4h (histological data) after music influences when the BBB integrity was fully restored, or 4 weeks after delayed music effects. We hypothesize that the listening of loud music in the repetitive mode can adapt the brain to sound stress and protect its structure from sound-induced injuries.

We have present in detail possible systemic and molecular mechanisms responsible for music-induced OBBB. There is the hypothesis that stress hormone, such as epinephrine, might induce an increase in the BBB permeability via vasodilation of cerebral vessels and an increase in the extension of cracks in the tight contacts of endothelial cells with a change in the ultrastructure of the astrocyte end-feet; increase transport and pinocytotic activity of endothelial cells [27–30]. The elevated level of stress hormones, including epinephrine, induced by sound, was demonstrated in humans and animals [47]. In our results we have shown that loud music induces a rise of serum epinephrine with a significant increase in CBF in both macro- and microcirculatory levels. We assume that elevation of epinephrine by stress causes an increase of CBF and changes in the tone of cerebral vessels that could be an initial factor triggering the BBB leakage. Indeed, 1h after music exposure, we have observed a decrease of the signal intensity from the TJ proteins, such as CLND5 and OCC, suggesting a temporal disorganization of the TJ assembly. It is important to note that already 4h after music effects, the signal intensity from the TJ proteins was within normal units that was preserved in the next day after the experiment. We hypothesize that these fast changes can be explained by the internalization of the TJ proteins resulting in a temporal loss of their surface in the space between endothelial cells that can be one of the mechanisms underlying music-induced OBBB. Our hypothesis is very close to the mechanism of BBB disruption via the internalization of VE-cadherin induced by VEGF [48]. Thus, the BBB breakdown induced by music exposure causes a reversible disorganization of the TJ machinery in affected cerebral endothelial cells.

Using a model of sensorineural deafness in mice, we have not found effect of sound on the BBB permeability to EBAC in mice with a hearing loss suggesting an important role of the auditory system in music-induced OBBB.

In sum, our findings clearly demonstrate that loud music induces a temporal OBBB with a fast recovery of BBB functions. The OBBB is accompanied by lymphatic clearance of molecules penetrating into the brain via OBBB that can be a crucial mechanism of quickly restoration of the BBB integrity after music exposure [38–40].

## Conclusion

This pioneering experimental study has discovered the important phenomenon of OBBB in healthy mouse brain induced by loud rock music (90-100 dB, 11 – 10.000 Hz, Scorpions “Still loving you”). The listening of loud music during 2h in intermittent adaptive mode is accompanied by delayed (1h after music exposure) and short-lasting (during 1-4 hrs) OBBB for high and low molecular weight molecules. We assume that an elevation of epinephrine by sound stress causes an increase of CBF and changes in the tone of cerebral vessels that could be an initial factor triggering the BBB leakage associated with a decrease in expression of TJ proteins. The music-induced OBBB is reversible with quickly restoration of the BBB function without the injuries of brain tissues and the cochlea. It can be related to the repetitive mode of music exposure and post-OBBB activation of lymphatic clearance of molecules penetrating into the brain via OBBB.

Our results stimulate a revision of the classic knowledge about the BBB nature. The BBB opens itself in a healthy brain as a response to sound stress. Despite the fact that we did not find any injuries of the brain tissues and the cochlea after loud music-OBBB due to limitation of our studies of delayed music effects by 4 weeks, even temporal OBBB can be harmful for the brain via opening the door for viruses, bacteria and toxins. It is important to note that millions of teenagers and professional rock, jazz and symphony orchestra musicians are in a potential risk group of OBBB due to exposure to loud music in concerts, nightclubs, discotheques, and pubs. Therefore, more detailed studies of loud music-induced OBBB are required for further investigations in human.

We also believe that our findings can be the basis for novel strategies in optimization of sound-mediated methods for brain drug delivery. Despite the fact that music opens BBB in a non targeted manner, loud music has a high potential for clinical applications as an easy used, non-invasive, low cost, labeling free, perspective and completely new approach for the treatment of neurodegenerative disorders, such as Alzheimer’s disease or myotrophic lateral sclerosis, which are associated with injuries of many brain regions and diffusive progression without specificity of the brain areas. We address to further research of music-induced OBBB as a proof of concept for exploration of sound-induced drug delivery to the brain. Such issues as the study of efficacy of the drug delivery into the brain and the effect of different levels of sound on the pharmacokinetics of the delivered drug can shed light on the development of a new alternative method of drug brain delivery for different neurodegenerative diseases, brain oncology and brain trauma.

## Material and Methods

### Subjects

The experiments were conducted on male mongrel mice (20-25 mg). All procedures were performed in accordance with the “Guide for the Care and Use of Laboratory Animals”. The experimental protocols were approved by the Local Bioethics Commission of the Saratov State University (Protocol No. 7) and the Institutional Animal Care and Use Committee of the University of New Mexico, USA (#200247).

**Experimental design of music effect on the BBB permeability** is presented in Figure 1 in SI To produce music (70-90-100 dB and 11-10,000 Hz, Scorpions “Still loving you”) we used loudspeaker (ranging of sound intensity – 0-130 dB, frequencies - 63-15000 Hz; 100 V, Yerasov Music Corporation, Saint Petersburg, Russia) (Figure 2 in SI). The repetitive music exposure was performed using the sequence of: 1 min – music on, then 1 min – music off during 2h. The sound level was measured directly in a cage of animals using the sound level meter (Megeon 92130, Russia) (Figure 3 in SI).

***In vivo* real time fluorescent microscopy of extravasation of EBAC** from the cerebral vessels into the brain tissues was performed via optical window [23] using the adapted protocol for two-photon imaging of the cortex in awake behavior mice [49] (See more details in SI).

### Spectrofluorometric assay of EBAC extravasation

The leakage of EBAC was determined in mice in four groups: I) control, no music, II, III and IV) 1, 4 and 24h after music exposure, respectively, n=15 in each group. Before or 1h/4h/24h after music interventions, Evans Blue dye (Sigma Chemical Co., St. Louis, Missouri, 2 mg⁄25 g mouse, 1% solution in physiological 0.9% saline) was injected into the femoral vein and circulated in the blood for 30 min. Then, the mice were decapitated, and their brains were quickly collected. To study the role of auditory system in music-BBBO, the EBAC leakage was evaluated in the intact and deafness mice, n=10 in each group. To exclude the effects of anesthesia on the BBB permeability, the EBAC level was evaluated in the additional group of mice received 2% isoflurane at 1L/min N_2_O/O_2_ – 70:30 during 30 min (the time of duration of anesthesia in *in vivo* experiments), n=10.

### Confocal microscopy of FITC-dextran 70 kDa extravasation

The confocal microscopy of the BBB permeability performed in the groups: 1) control, no music; 2) 1h; 3) 4h; 4) 24h after music exposure; n=10 in each group. Before or 1h/4h/24h after music interventions, Fluorescein isothiocyanate (FITC)-dextran 70 kDa (FTICD) (1 mg/25 g mouse, 0.5% solution in saline, Sigma-Aldrich) was injected into the tail vein and allowed to circulate for 2 min. Afterward, mice were decapitated and the brains were quickly removed and fixed in 4% paraformaldehyde (PFA) for 24 h, cut into 50-μm thick slices on a vibratome (Leica VT 1000S Microsystem, Germany) and analyzed using a confocal microscope (Olympus FV10i-W, Olympus, Japan).

### In vivo real-time two-photon laser scanning microscopy (2PLSM)

The BBB permeability via optical window [23] was continuously monitored by measuring the perivascular tissue fluorescence of FITCD 70 kDa (Sigma-Aldrich, in saline 5% wt/vol) in 10 mice at different time points: before or 1, 4 and 24h after music exposure as described previously with some modifications [50]. During the imaging mice were kept under inhalation anesthesia with 2% isoflurane at 1L/min N_2_O/O_2_ – 70:30. FITCD was injected through the tail vein (~100 μl) at an estimated initial concentration in blood serum of 150 μM. The BBB permeability was evaluated by measuring changes in perivascular tissue fluorescence in planar images of the cortex taken 50 and 150 μm depth in 20 min after FITCD injection using Olympus microscope (Japan) [50].

### MRI analysis of the BBB permeability

The MRI with gadolinium-diethylene-triamine-pentaacetic acid (Gd-DTPA, MW= 938 Da; Bayer Healthcare, 0.1 mM/kg) was conducted on the same mice, which we used for 2PLSM, at different time intervals 0 – before and 1, 4, 24h after sound exposure on a 7-T dedicated research MRI scanner (Bruker Biospin; Billerica, MA, USA). Signal transmission and detection was done with a small-bore linear RF coil (inner diameter of 72 mm) and a single tuned surface coil (RAPID Biomedical, Rimpar, Germany). The mice kept under inhalation anesthesia (2% isoflurane at 1L/min N2O/O2 – 70:30). To non-invasively evaluate the BBB permeability, we used a modified dynamic contrast-enhanced (DCE)-MRI and graphical analysis of the resultant image data [51].

### Biochemical assays of plasma epinephrine

The plasma epinephrine level (ng/ml) was determined using ELISA kits (Abnova, Taiwan) before music, immediately, 1h and 4h after music exposure in mice (n=10 in each group). The plates were read at 450 nm using an ELx 800 plate reader (BioTek Instruments Inc.). The detection limits were 0.3 ng/ml (with intra and inter assay variation coefficients of 11.2-16.3% and 8.7-12.6%, respectively).

### Laser speckle contrast imaging of rCBF

A custom-made laser speckle contrast imaging system was used to monitor rCBF before, immediately and 1h/4h/24h after sound exposure in 10 mice under the inhalation anesthesia (2% isoflurane, 70% N_2_O and 30% O_2_) via an optically cleared skull window (diameter: 5 mm) using optical clearing method FDISCO described in detail in Ref. [23]. The automated segmentation algorithm, described in our previous work [52], was used to calculate the mean value of CBF at macro- (in the Sagittal sinus) and micro-levels.

### Immunohistochemical assay

Mice in the control group (before music, n=10) and in experimental groups immediately, 1-4-24 hrs after music exposure (n=10 in each group) were euthanized with an intraperitoneal injection of a lethal dose of ketamine and xylazine and intracardially perfused with 0.1 M of PBS for 5 min. Afterward, the brains were removed and fixed in 4% buffered paraformaldehyde for one day and in 20% sucrose for another day. The signal intensity from the examined proteins were evaluated on free-floating sections using the standard method of simultaneously combined staining (Abcam Protocol). Brain slices (50 μm) were blocked in 150 μl 10% BSA/0.2% Triton X-100/PBS for 2 h, then incubated overnight at 4 C and 2 h at room temperature with CLND-5, ZO-1 (1:500; Santa Cruz Biotechnology, Santa Cruz, USA), OCC and NG2 (1:500; Abcam, Cambridge, UK). After several rinses in PBS, the slides were incubated for 3h at room temperature with fluorescent- labeled secondary antibodies on 1% BSA/0.2% Triton X-100 /PBS (1:500; Goat A/Rb, Alexa 555 and 647 Abcam, UK). Confocal microscopy of the cerebral cortex was performed using confocal microscope with water immersion Olympus FV10i-W (Olympus, Japan). In all cases, 10 regions of interest were analyzed.

### Assessment of apoptosis with TUNEL method

The number of apoptotic cells before and 1h and 4 weeks after music exposure (n=10 in each group) was evaluated with the TUNEL method using the “17- 141 TUNEL Apoptosis Detection Kit” (Abcam, UK) in accordance with the standard protocol provided by the manufacturer and analyzed by confocal microscopy (Olympus FLUOVIEW FV10i-W, Tokyo, Japan).

For **histological analysis of the brain tissues and SGCs** (see details of embedding of auditory bulla and capsule in SI), four groups of animals were used: I) control group, no music; II, III and IV – experimental groups, 1h and 4 weeks after music exposure, respectively; n=10 in each group. All mice were euthanized with an intraperitoneal injection of a lethal dose of ketamine and xylazine. Afterward, the brains were removed and fixed in 10% buffered paraformaldehyde. The paraformaldehyde-fixed specimens were embedded in paraffin, sectioned (4 μm) and stained with hematoxylin and eosin. The histological sections were evaluated by light microscopy using the digital image analysis system Mikrovizor medical μVizo-103(LOMO,Russia).

### A model of sensorineural deafness

To establish animal model of deafness, we used a synergistic ototoxic effect of single administration of Furosemide (100 mg/kg, iv, St. Louis, MO, USA) and Kanamycin sulfate (1000 mg/kg, im, St. Louis, MO, USA) [53,54] (See more details in SI).

### The study of lymphatic clearance of FITCD from the brain after OBBB

1 h after music-OBBB, FITCD was injected intravenously (1 mg/25 g mouse, 0.5% solution in 0.9% physiological saline, Sigma-Aldrich, St. Louis, USA) and allowed to circulate for 30 min. Afterward, mice were decapitated; their dcLNs were removed and fixed in 4% buffered PFA for one day. To label the lymphatic and blood vessels, samples were incubated overnight at +4C with goat anti-rabbit Lyve-1 and Prox-1 antibody (1:500; Invitrogen, Molecular Probes, Eugene, Oregon, USA). After several rinses in PBS, the samples were incubated for 3h at room temperature with fluorescent-labeled secondary antibodies on 1% BSA/0.2% Triton X-100 /PBS (1:500; goat anti-rabbit IgG (H+L) Alexa Four 555 and goat anti-mouse IgG (H+L) Alexa Four 647; Invitrogen, Molecular Probes, Eugene, Oregon, USA) with further confocal analysis (Olympus FV10i-W, Olympus, Japan).

### Statistical analysis

The results are presented as mean ± standard error of the mean (SEM). Differences from the initial level in the same group were evaluated by the Wilcoxon test. Intergroup differences were evaluated using the Mann-Whitney test and ANOVA-2 (post hoc analysis with Duncan’s rank test). The significance levels were set at p < 0.05-0.001 for all analyses.

## Acknowledgments

We would like to express our special thanks of gratitude to Prof. John Connor (University of New Mexico) for discussion our results and many helpful advices for preparation of the manuscript.

## Finding

S-G O., AS, NN, JK were supported by RF Governmental Grant № 075-5-2019-885, Grant from RSF № 20-5-00090 and 19-5-00201, Grant from RFBR 19-515-55016 China a, 20-015-00308-a. DB was supported by NIH R21NS091600 and P20GM109089.

## Notes

https://drive.google.com/file/d/1Ee-Z2QIC7m0tq1a4LQJQhdmTbIoPiMsr/view?usp=sharing

https://youtu.be/CMczv1RYWtw

https://youtu.be/gcvMWYAzWoA

https://youtu.be/UIpaQOAAesA

https://youtu.be/BuXz-NG9lNQ

### Competing Interest Statement

The authors have declared no competing interest.

## References

1. Koelsch S. 2014 Brain correlates of music-evoked emotions. Nat Rev Neurosci., 15(3), 170–180. (doi: 10.1038/nrn3666)

2. Mitterschiffthaler MT, Fu CH, Dalton JA, Andrew CM, Williams SC. 2007 A functional MRI study of happy and sad affective states induced by classical music. Hum Brain Mapp., 28(11), 1150–1162. (doi: 10.1002/hbm.20337)

3. Music and the Brain. Edited by M. Critchley and R. A. Henson. (Pp. 458; illustrated; £11.50.) Heinemann: London. 1977. Volume 7 Issue 4.

4. Stewart L., Kriegstein K., Warren J.D., Griffiths T.D. 2006 Music and the brain: disorders of musical listening. Brain, 129, 2533–2553. (doi.org/10.1093/brain/awl171)

5. WHO (World Health Organization) (2020). Make Listening Safe. Available online at: (http://www.who.int/pbd/deafness/activities/MLS)

6. Yassi A, Pollock N, Tran N, Cheang M. 1993 Risks to hearing from a rock concert. Can Fam Physician, 39, 1045–50.

7. Opperman D.A., Reifman W, Schlauch R, Levine S. 2006 Incidence of spontaneous hearing threshold shifts during modern concert performances. Otolaryngol Head Neck Surg, 134(4), 667–673. (doi: 10.1016/j.otohns.2005.11.039)

8. Clark WW. 1991 Noise exposure from leisure activities: A review. J Acoust Soc Am, 90(1), 175–178. (doi: 10.1121/1.401285)

9. Canadian Centre for Occupational Health and Safety [homepage on the Internet]. Hamilton: The Centre; c2007. [updated 2007 Mar 19; cited 2007 May 20]. Available from: http://www.ccohs.ca/oshanswers/phys_agents/exposure_can.html

10. American Academy of Audiology [homepage on the Internet]. Reston: The Academy; c2007 [updated 2003; cited 2007 May 20]. Available from: www.audiology.org/publications/documents/positions/Hearingconservation.

11. Stormer C.C, Stenklev N.C. 2007 Rock music and Hearing Disorders. Tidsskr Nor Laegeforen, 127(7), 874–877. (doi: 10.1121/1.401285)

12. Kahari K, Zachau G, Eklof M, Sandsjo L, Moller C. 2003 Assessment of hearing and hearing disorders in rock/jazz musicians. Int J Audiol, 42, 279–288. (doi: 10.3109/14992020309078347)

13. Teie PU. 1998 Noise-induced hearing loss and symphony orchestra musicians: risk, factors, effects, and management. Md Med J, 47(1), 13–8.

14. Petrescu N. 2008 Loud Music Listening. MJM, 11(2), 169–176.

15. Shi X. 2016 Pathophysiology of the cochlear intrastrial fluid-blood barrier. Hear Res, 338, 52–63. (doi:10.1016/j.heares.2016.01.010)

16. Inamura N, Salt A.N. 1993 Permeability changes of the blood-labyrinth barrier measured in vivo during experimental treatments. Hear Res, 61(1-2), 12–18. (doi.org/10.1016/0378-5955(92)90030-Q)

17. Shi X. 2009 Cochlear Pericyte Responses to Acoustic Trauma and the Involvement of Hypoxia-Inducible Factor-1α and Vascular Endothelial Growth Factor. The American Journal of Pathology, 174(5), 1692–1704. (doi: 10.2353/ajpath.2009.080739)

18. Shi X, Nuttall AL. 2007: Expression of adhesion molecular proteins in the cochlear lateral wall of normal and PARP-1 mutant mice. Hear Res, 224, 1–14. (doi: 10.1016/j.heares.2006.10.011)

19. Abbott J, Patabendige A, Dolman D, Yusof S, Begley D. 2010 Structure and function of the blood–brain barrier. Neurobiology of Disease, 37, 13–25. (doi.org/10.1016/j.nbd.2009.07.030)

20. http://www.asha.org/public/hearing/Noise/

21. http://www.industrialnoisecontrol.com/comparative-noise-examples.htm

22. Namykin A, Shushunova N, Ulanova M, Semyachkina-Glushkovskaya O, Tuchin V, Fedosov I. 2018 Intravital molecular tagging velocimetry of cerebral blood flow using Evans Blue. J. Biophotonics, 11(8): e201700343. (doi: 10.1002/jbio.201700343)

23. Yisong Qi, Tingting Yu, Jianyi Xu, et. al. 2019 FDISCO: Advanced solvent-based clearing method for imaging whole organs. Sci. Adv, 5: eaau8355. (doi: 10.1126/sciadv.aau8355)

24. https://www.hear-it.org/disco-noise-volume-over-the-top-1

25. Saunders N, Dziegielewska K, Møllgård K, Habgood M. 2015 Markers for blood-brain barrier integrity: how appropriate is Evans blue in the twenty-first century and what are the alternatives? Front. Neurosci, 9:385. (doi: 10.3389/fnins.2015.00385)

26. Yang E-I, Lin E.W., Hensch T.K. 2012 Critical period for acoustic preference in mice. PNAS, 109(Supplement 2) 17213–17220. (doi:10.1073/pnas.1200705109)

27. Akihiko U, Grubb J, Banks W, Sly W. 2007 Epinephrine enhances lyposomal enzyme delivery across the blood-brain barrier by up-regulation of the mannose 6-phosphate receptor. Proc. Natl. Acad. Sci. USA, 104(31):12873–8. (doi: 10.1073/pnas.0705611104)

28. Johansson B, Martinsson L. 1980 The blood-brain barrier in adrenaline-induced hypertension: circadian variations and modification by beta-adrenoreceptor antagonists. Acta. Neurol. Scand, 62(2):96–102. (doi: 10.1111/j.1600-0404.1980.tb03009.x)

29. Murphy V, Johanson C. 1985 Adrenergic-induced enhancement of brain barrier system permebility to small nonelectrolyes: choroid plexus versus cerebral capillaries. J. Cereb. Blood Flow Metab, 5(3):401–12. (doi: 10.1038/jcbfm.1985.55)

30. Sarmento A, Borges N, Azevedo I. 1991 Adrenergic influences on the control of blood-brain barrier permeability. Naunyn-Schmiedeberg’s Archives of Pharmacology. 343:633–637. (doi.org/10.1007/BF00184295)

31. Liberman M.C., Kujawa S.G. 2017 Cochlear synaptopathy in acquired sensorineural hearing loss: Manifestations and mechanisms. Hear Res. 349, 138–147. (doi:10.1016/j.heares.2017.01.003)

32. Lawner B.E., Harding G.W., Bohne B.A. 1997 Time course of nerve-fiber regeneration in the noise-damaged mammalian cochlea. Int J Dev Neurosci, 15(4–5), 601–617. (doi.org/10.1016/S0736-5748(96)00115-3)

33. Wang Y, Hirose K, Liberman M.C. 2002 Dynamics of noise-induced cellular injury and repair in the mouse cochlea. J Assoc Res Otolaryngol. 3(3):248–268. (doi: 10.1007/s101620020028)

34. Webster D.B., Webster M. 1978 Cochlear nerve projections following organ of corti destruction. Otolaryngol. 86(2):342–353. (doi.org/10.1177/019459987808600228)

35. Sugawara M, Corfas G, Liberman M.C. 2005 Influence of supporting cells on neuronal degeneration after hair cell loss. J Assoc Res Otolaryngol, 6(2):136–147. (doi: 10.1007/s10162-004-5050-1)

36. McDannold N, Vykhodtseva N, Raymond S, Jolesz F.A, Hynynen K. 2005 MRI-guided targeted blood-brain barrier disruption with focused ultrasound: histological findings in rabbits. Ultrasound Med Biol, 31(11):1527–1537. (doi: 10.1016/j.ultrasmedbio.2005.07.010)

37. Hynynen K, McDannold N, Vykhodtseva N, Raymond S, Weissleder R, Jolesz F.A., Sheikov N.S. 2006 Focal disruption of the blood–brain barrier due to 260-kHz ultrasound bursts: a method for molecular imaging and targeted drug delivery. J Neurosurg, 105(3):445–454. (doi: 10.3171/jns.2006.105.3.445)

38. Semyachkina-Glushkovskaya O., Chehonin V, Borisova E. et al. 2018 Photodynamic opening of the blood-brain barrier and pathways of brain clearing pathways. J Biophotonics, 11(8):e201700287. (doi: 10.1002/jbio.201700287)

39. Semyachkina-Glushkovskaya O., Abdurashitov A., Dubrovsky A. et al. 2017 Application of optical coherent tomography for in vivo monitoring of the meningeal lymphatic vessels during opening of blood-brain barrier: mechanisms of brain clearing. JBO, 22(12):1–9. (doi: 10.1117/1.JBO.22.12.121719)

40. Semyachkina-Glushkovskaya O, Bragin D, Bragina O, Yang Y, Abdurashitov A, Esmat A. et al. Mechanisms of sound-induced opening of the blood-brain barrier. 2020 Adv. Exp. Med. Biol, 1269(in press).

41. Semyachkina-Glushkovskaya O., Postnov D., Kurths J. 2018 Blood–Brain, Barrier, Lymphatic, Clearance, and Recovery: Ariadne’s Thread in Labyrinths of Hypotheses. Int. J. Mol. Sci., 19(12), 3818. (doi.org/10.3390/ijms19123818)

42. Reynolds R.P., Kinard W.L., Degraff J.J., Leverage N., Norton J.N. 2010 Noise in a Laboratory Animal Facility from the Human and Mouse Perspectives. JAALAS, 49(5), 592–597.

43. Grill-Spector K, Henson R, Martin A. 2006 Repetition and the brain: neural models of stimulus-specific effects. Trends Cogn Sci, 10(1):14–23. (doi.org/10.1016/j.tics.2005.11.006)

44. Sobotka S., Ringo J.L. 1994 Stimulus specific adaptation in excited but not in inhibited cells in inferotemporal cortex of macaque. Brain Res, 646, 95–99. (doi.org/10.1016/0006-8993(94)90061-2)

45. Ringo J.L. 1996 Stimulus specific adaptation in inferior temporal and medial temporal cortex of the monkey. Behav. Brain Res, 76, 191–197. (doi.org/10.1016/0166-4328(95)00197-2)

46. Grill-Spector K. and, Malach R. 2001 fMR-adaptation: a tool for studying the functional properties of human cortical neurons. Acta Psychol. (Amst.), 107, 293–321. (doi.org/10.1016/S0001-6918(01)00019-1)

47. Turner J, Parrish J, Hughes L, Toth L, Caspary D. 2005 Hearing in laboratory animals: strain differences and nonauditory effects of noise. Comp Med, 55(1), 12–23. (https://www.ncbi.nlm.nih.gov/pmc/articles/PMC3725606)

48. Hebba J, Lecrair H, Azzi S, Roussel C, Scott M, Bidere N, Gavard J. 2013 The C-terminus region of β-arrestin-1 modulates VE-cadherin expression and endothelial cell permeability. J. Cell. Commun. Signal, 11,37. (doi: 10.1186/1478-811X-11-37)

49. Villette V., Chavarha M., Dimov I. et al. Ultrafast Two-Photon Imaging of a High-Gain Voltage Indicator in Awake Behaving Mice. Cell, 179(7), 1590–1608.e23. (doi.org/10.1016/j.cell.2019.11.004)

50. Bragin D, Kameneva M, Bragina O, Thomson S, Statom G, Lara D, Yang Y, Nemoto E. 2017 Rheological effects of drug-reducing polymers improve cerebral blood flow and oxygenation after traumatic brain injury in rats. J. Cereb. Blood Flow Metab. 37(3), 762–775. (doi: 10.1177/0271678X16684153)

51. Patlak S.C, Blasberg R.G, Fenstermacher J.D. 1983 Graphical evaluation of blood-to-brain transfer constants from multiple-time uptake data. J. Cereb. Blood Flow Metab. 3:1–7. (doi: 10.1038/jcbfm.1983.1)

52. Abdurashitov A, Lychagov V, Sindeeva O, Semyachkina-Glushkovskaya O, Tuchin V. 2015 Histogram analysis of laser speckle contrast image for cerebral blood flow monitoring. Front. Optoelectron. 8(2), 187–194. (doi.org/10.1007/s12200-015-0493-z)

53. Long M, Hai-jin Y, Fen-qian Y, Wei-wei G, Shi-ming Y. 2015 An e cient strategy for establishing a model of sensorineural deafness in rats. Neural Regen Res, 10(10): 1683–1689. (doi: 10.4103/1673-5374.153704)

54. Liberman M, Gao J, He D, Wu X, Jia S, Zuo J. 2002 Prestin is required for electromotility of the outer hair cell and for the cochlear amplifier. Nature, 419:300–304. (doi: 10.1038/nature01059)

