## Supplemental material for "Phenomenon of music-induced opening of the blood-brain barrier in healthy mice"

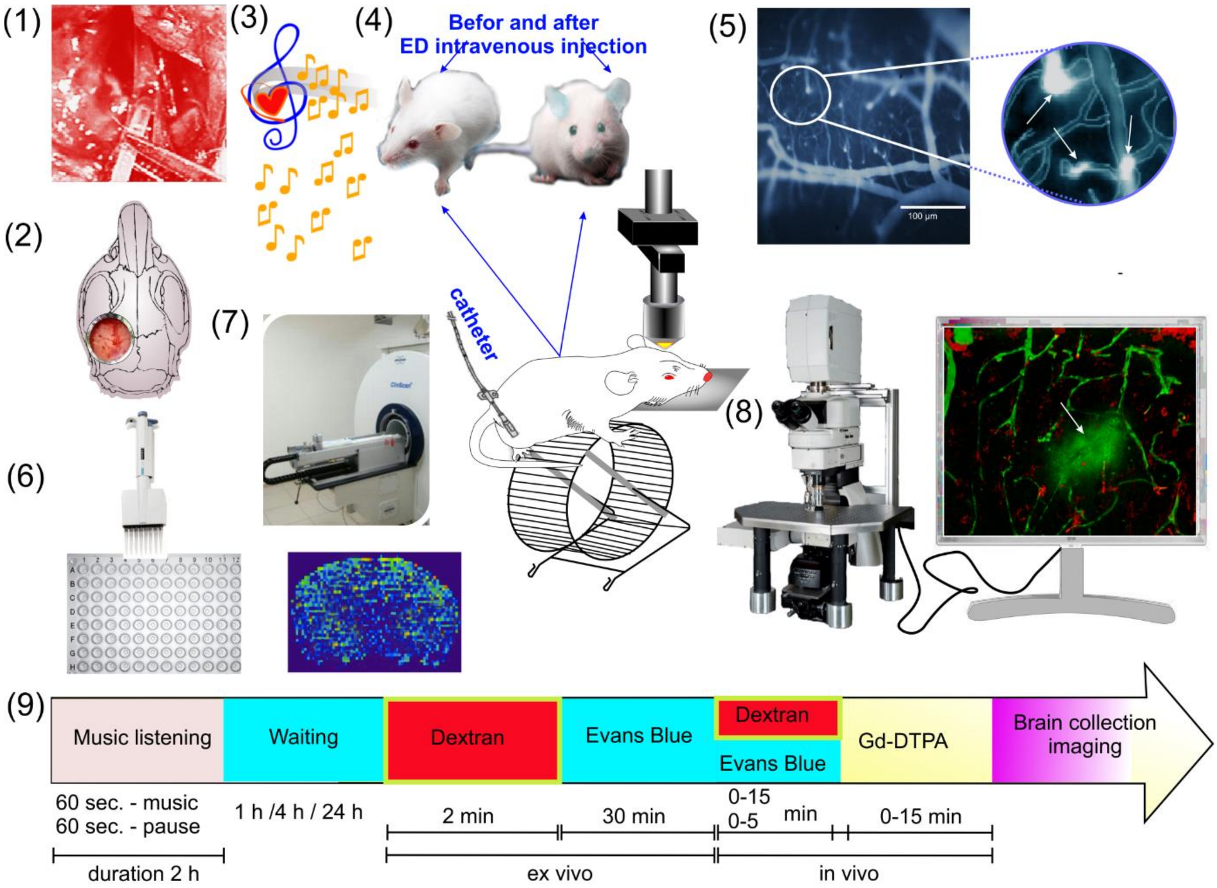


**Fig. 1 SI. Design of *in vivo* and *ex vivo* experiments of the study of loud music-induced opening of the blood-brain barrier (OBBB)**: (1) three days before experiments the polyethylene catheter (PE-10 tip, Scientific Commodities Inc., Lake Havasu City, Arizona) was implanted into the femoral vein for injection of tracers (Evans Blue dye, fluorescein isothiocyanate dextran 70 kDa, gadolinium-diethylene-triamine-pentaacetic acid - Gd-DTPA); (2) – the optical window [1] was prepared for *in vivo* real time fluorescent microscopy of the BBB permeability to the Evans Blue Albumin Complex (EBAC, 68.5 kDa) in awake behavior mice (see session "In vivo real time fluorescent microscopy of extravasation of Evans Blue"); (3) – afterward mice (n=15 in each group) were underwent to the intermittent music (70-90-100 dB, 11-10,000 Hz, Scorpions "Still loving you") during 2h (60 sec sound and 60 sec – pause) (see session "Experimental design of music effect on the BBB permeability"); (4) – Evans Blue dye was injected via catheter immediately after music-off and then we performed *in vivo* real time fluorescent microscopy of OBBB for EBAC in awake behavior mice during 5 hrs; (5) The EBAC leakage was detected as bright fluorescence around the cerebral microvessels; (6) after *in vivo* real time fluorescent microscopy of OBBB for EBAC, all mice were decapitated, their brain removed and analyzed using spectrofluorometric assay of EBAC extravasation (see session "Spectrofluorometric assay of EBAC extravasation"); (7) additionally, magnetic resonance imaging (MRI) was used for the study of the BBB permeability for Gd-DTPA; (8) also *ex vivo* confocal imaging and *in vivo* real-time two-photon laser scanning microscopy of FITC-dextran extravasation was performed; (9) the time points for injections of the model compounds in relation to the music intervention and the brain collection.

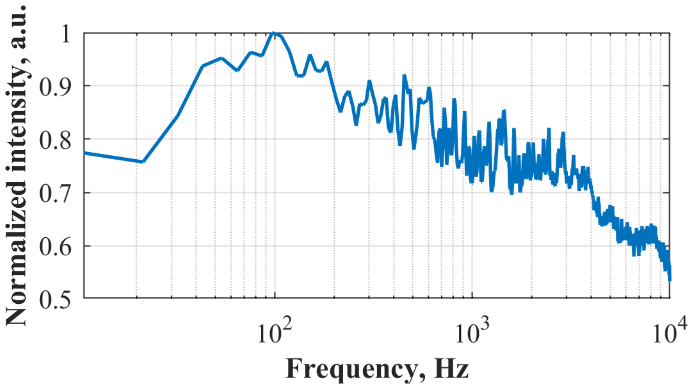


**Fig. 2 SI. The frequency range of music (Scorpions “Still Loving You”)**: frequencies in the range of 11-10,000 Hz and maximal intensity around 100 Hz.

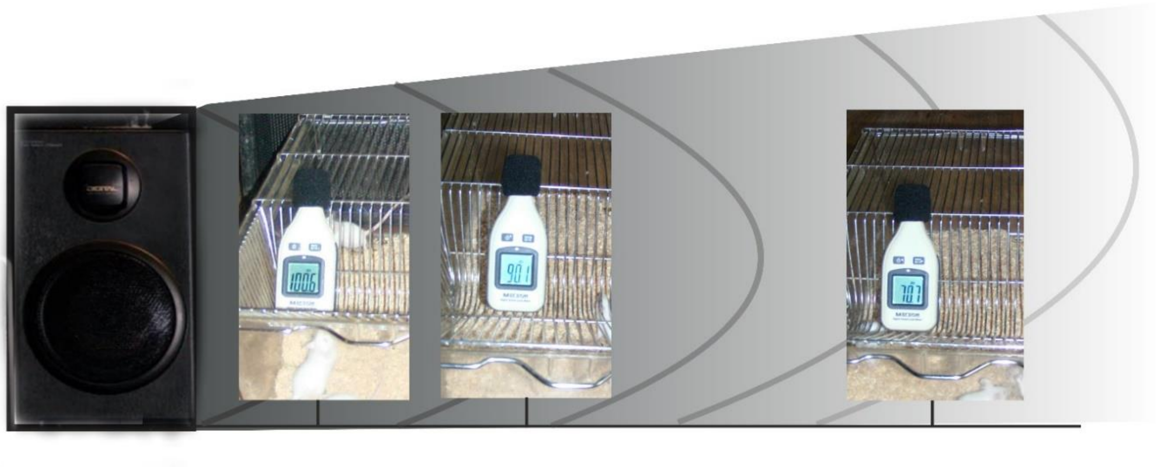

**Fig. 3 SI. The measure of sound level in the cages of animals during listening of loud music.** To produce the loud music, we used loudspeaker generated sound 100 dB. The location of cage with mice from loudspeaker was: 1) at 1m - 100 dB; 2) at 2m - 90 dB; 3) at 32m - 70 dB in according with information referred here [https://lgmproducts.com/technical-information/sound-pressure-levels-in-dba]. The sound energy was measured directly in a cage of animals using sound level meter (Megeon 92130, Russia).

### Table 1 SI. The effect of loud music on the BBB permeability to EBAC (µg/g tissue)

| **Sound level (dB) and time elapsed after sound exposure (h)** | **Content of EBAC (µg/g tissue)** | | |
|---|---|---|---|
| No music (the control group) | 0.15±0.01 | | |
| | **Music** duration during 2 h (intermittent) | **Music** duration during 1 h (intermittent) | **Music** duration during 0.25 h (continues) |
| 100 dB | | | |
| immediately | 0.12±0.01 | 0.11±0.05 | 0.14±0.02 |
| 1 h | **2.60±0.06** *** | 0.12±0.03 | 0.17±0.01 |
| 4 h | 0.19±0.03 | 0.15±0.07 | 0.15±0.05 |
| 24 h | 0.16±0.03 | 0.11±0.09 | 0.17±0.02 |
| 90 dB | | | |
| immediately | 0.15±0.08 | 0.11±0.06 | 0.17±0.01 |
| 1 h | **2.70±0.04** ***(n=11) | 0.16±0.03 | 0.16±0.08 |
| 4 h | 0.18±0.06 (n=4)# | 0.14±0.03 | 0.14±0.05 |
| 24 h | 0.15±0.03 | 0.17±0.02 | 0.18±0.01 |
| | 0.19±0.07 | | |
| 70 dB | | | |
| immediately | 0.13±0.02 | 0.16±0.07 | 0.12±0.03 |
| 1 h | 0.17±0.08 | 0.11±0.04 | 0.14±0.09 |
| 4 h | 0.19±0.06 | 0.13±0.02 | 0.14±0.02 |
| 24 h | 0.19±0.09 | 0.18±0.02 | 0.11±0.01 |

$p<0.001$: *** - vs. before music exposure (the control group), n=15 for the groups (music duration 2h) and n=10 for the groups (music duration 0.25-1 h); # - the number of mice without the BBB opening.

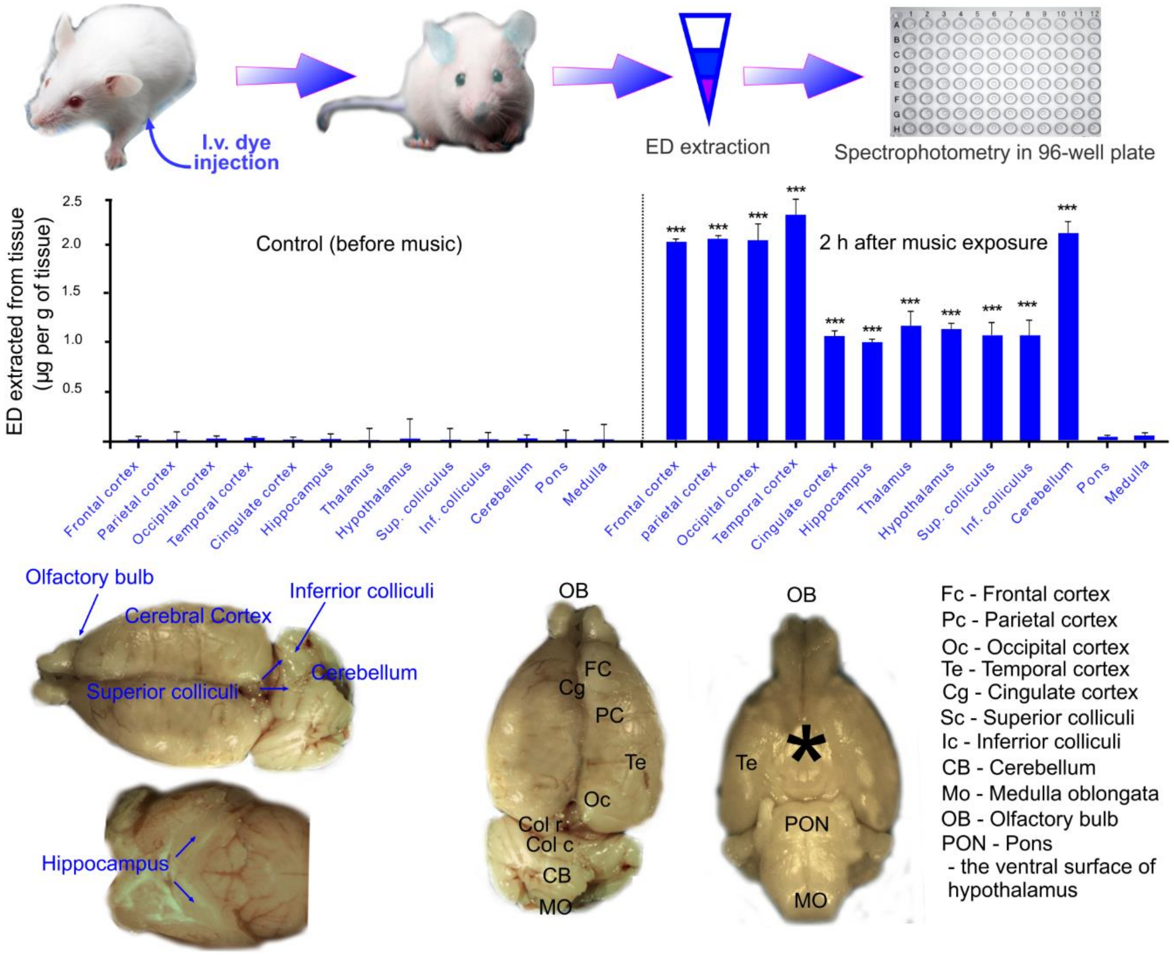


**Fig. 4 SI. The effect of loud music on the BBB permeability to EBAC (µg/g tissue) in different brain fields**: *** - $p<0.001$ vs. the control group (no music), n=15 in each group.

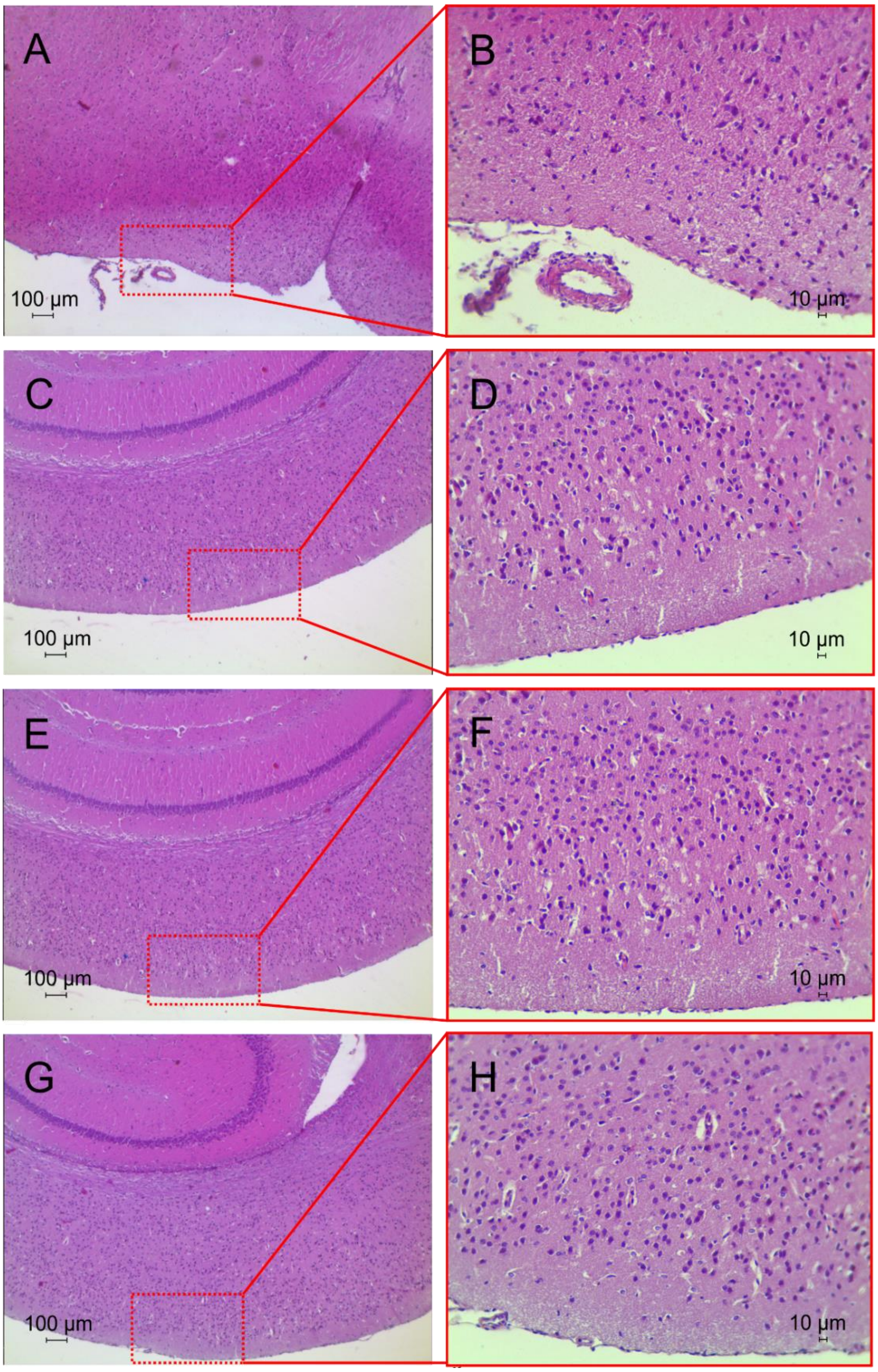


**Fig. 5 SI. Histological analysis of the brain tissues before and after loud music exposure**: A and B – the control group, no music influences, C and D, E and F, G and H – 1h, 4h and 4 weeks after music exposure, respectively, n=10 in each group. Hematoxylin & Eosin staining. Bars represent 10 µm (246.4X).

**Video resources:**

Video 1 - https://youtu.be/CMczv1RYWtw

Video 2 – https://youtu.be/gcvMWYAzWoA

Video 3 – https://youtu.be/UIpaQOAAesA

Video 4 – https://youtu.be/BuXz-NG9lNQ

### Additional details of methods used in experiments

***In vivo* real time fluorescent microscopy of extravasation of Evans Blue** from the cerebral vessels into the brain tissues was performed via optical window using adapted protocol for two-photon imaging of the cortex in awake behavior mice [2]. Fifteen min before imaging, optical window [1] was prepared in coordinates 1-4 mm caudal and 1-4 mm lateral to bregma. Preliminary 3 days before experiment, a polyethylene catheter (PE-10 tip, Scientific Commodities Inc., Lake Havasu City, Arizona) was inserted into the right femoral vein for Evans Blue intravenous injection in a single bolus dose (2 mg⁄100 g, 1% solution in physiological 0.9% saline). Microscope (Axio Imager A1, Zeiss, Germany) was equipped with CMOS camera (acA1920-40uc, Basler AG, Germany), 10× 0.3 objective lens and Evans Blue dye filter sets 49019 (Chroma, USA). Continuous wave laser diode module (50000463, Laserlands.net, China) with 160 mW output power at 635 nm was used to excite the dye florescence. Laser beam was expanded with a cylindrical lens (f = 50 mm) and then directed towards an object at 45° with respect to the microscope optical axis. To reduce the sample irradiation, the laser was synchronized with the camera "fire" output to turn it on only for the image capturing period. The awake behavior mice were positioned at the microscope stage using 3D printed homemade system for moving paws in a rotating circle and fixation of head. To adapt mice for experimental condition, they trained to be with fixed head and move paws on rotating sphere in the microscope system without any performance during two weeks (Figure 1 in SI).

**A model of sensorineural deafness**.

To establish animal model of deafness, we used a synergistic ototoxic effect of single administration of Furosemide (100 mg/kg, iv, St. Louis, MO, USA) and Kanamycin sulfate (1000 mg/kg, im, St. Louis, MO, USA) [53,54]. Auditory brainstem response measurements for confirmation of stable long-term hearing loss were performed 3 days after drugs administration as described previously [3]. Briefly, the sculp electrodes were inserted at the vertex and pinna in anesthetized mice (xylazine, 0.1 mg/kg and ketamine, 30 mg/kg). A series of 5-ms tone pips and clicks were presented at a high rate of speed. Levels were incremented in 5 dB steps from 10 to 100 dB SPL. Each click evoked waves of neural activity in the brainstem that were computer-averaged so they are differentiated from non-auditory background voltages. Both ears were measured.

**Embedding of auditory bulla and capsule in paraffin blocks** was performed using protocol published in Ref. 4. Mice were euthanized with an intraperitoneal injection of a lethal dose of ketamine and xylazine. Then head was decapitated and the skin was peeled completely towards the nose and cut off together with the snout and incisors. Then scissors were inserted into the mouth and cut masseter muscles on both sides. The jaw was opened carefully and removed together with the tongue. Using sharp scissors, the skull was skilled into two halves along the midsagittal plane. The cerebral and cerebellar hemispheres together with the brainstem were removed. Under a binocular microscope, the bulla and capsule with the surrounding skull bone were dissected. The anterior end of the bulla was cut with scissors and allowed 4% paraformaldehyde (PFA) in PBS to enter into the bulla. Then the bulla and capsule left in the fixative at 4 °C O/N on a tube rotator. Decalcification of the bulla and the capsule was made for a week at 4 °C in 10% ethylenediaminetetraacetic acid disodium salt dihydrate (EDTA-2Na), 100 mM Tris base, pH 7.0, in a 2 mL tube. The buffer was changed every other day. Afterward, the samples were removed and fixed in 4% buffered PFA. The paraformaldehyde-fixed specimens were embedded in paraffin, sectioned (4 μm) and stained with hematoxylin and eosin. The histological sections were evaluated by light microscopy using the digital image analysis system Mikrovizor medical μVizo-103(LOMO, Russia).

### References:


1. Yisong Qi, Tingting Yu, Jianyi Xu et. al. 2019 FDISCO: Advanced solvent-based clearing method for imaging whole organs. Sci. Adv, 5: eaau8355. (doi: 10.1126/sciadv.aau8355)
2. Villette V., Chavarha M., Dimov I. et al. Ultrafast Two-Photon Imaging of a High-Gain Voltage Indicator in Awake Behaving Mice. Cell, 179(7), 1590-1608.e23. (doi.org/10.1016/j.cell.2019.11.004)
3. Liberman M, Gao J, He D, Wu X, Jia S, Zuo J. 2002 Prestin is required for electromotility of the outer hair cell and for the cochlear amplifier. Nature, 419:300–304. (doi: 10.1038/nature01059)
4. Sakamoto, A., Kuroda, Y., Kanzaki, S., Matsuo, K. 2017 Dissection of the Auditory Bulla in Postnatal Mice: Isolation of the Middle Ear Bones and Histological Analysis. *J. Vis. Exp*. **11**9, e55054. (doi:10.3791/55054)
